# Transient Hsp90 suppression promotes a heritable change in protein translation

**DOI:** 10.1101/366070

**Authors:** Peter Tsvetkov, Zarina Brune, Timothy J. Eisen, Sven Heinrich, Greg A. Newby, Erinc Hallacli, Can Kayatekin, David Pincus, Susan Lindquist

**Affiliations:** Whitehead Institute for Biomedical Research, Cambridge, MA 02142; Massachusetts Institute of Technology, Cambridge, MA 02142, USA; Howard Hughes Medical Institute, Cambridge, MA 02139, USA

**Author notes:** Deceased (Susan Lindquist).

## Abstract

The heat shock protein 90 (Hsp90) chaperone functions as a protein-folding buffer and plays a unique role promoting the evolution of new heritable traits. To investigate the role of Hsp90 in modulating protein synthesis, we screened more than 1200 proteins involved in mRNA regulation for physical interactions with Hsp90 in human cells. Among the top hits was CPEB2, which strongly binds Hsp90 via its prion domain, reminiscent of the prion-like regulation of translation of *Aplysia* CPEB. In a yeast model of CPEB prion-dependent translation regulation, transient inhibition of Hsp90 amplified CPEB2 prion activity and resulted in persistent translation of the CPEB reporter. Remarkably, inhibition of Hsp90 was sufficient to induce a heritable change in protein translation that persisted for 30 generations, even in the absence of exogenous CPEB. Although we identified a variety of perturbations that enhanced translation of the reporter, only Hsp90 inhibition led to persistent activation. Thus, transient loss of Hsp90 function leads to the non-genetic inheritance of a novel translational state. We propose that, in addition to sculpting the conformational landscape of the proteome, Hsp90 promotes phenotypic variation by modulating protein synthesis.

## Introduction

Across evolution many of the molecular pathways underlying complex multicellular processes such as development and neuronal synapse formation evolved from basic environmental signaling circuits in unicellular organisms like yeast [1, 2]. The regulation of protein translation is a key hub that integrates extracellular signals into a cellular response [3, 4]. Indeed, the requirement for specific translation regulating factors to integrate environmental cues and induce specific mRNA translation is critical in development, stress response and neuronal function [4, 5], and also for the survival of single-celled organisms.

The members of the cytoplasmic polyadenylating element binding (CPEB) family of RNA-binding proteins act as regulators of development [6, 7] and synaptic protein synthesis [8, 9], aiding in synaptic remodeling and the persistence of behavioral memories [10, 11]. The CPEB homologs can be divided into two evolutionarily conserved groups. The first group includes human CPEB1 and its orthologs from *Xenopus* and *Drosophila (Orb1)* [12]. The second group includes human CPEB2-4, the neuronal *Aplysia* CPEB and the *Drosophila* ortholog, Orb2, all of which have been shown to regulate synaptic function. This group of CPEBs possess a glutamine-rich N-terminal domain thought to comprise a functional prion domain (PrD). The ability of neuronal CPEBs to activate translation in response to synaptic activity depends largely on the prion domain [9, 13-19]; in the aggregated prion form, the neuronal CPEB is active. In the soluble form, it is dormant [13, 19]. The prion properties of *Aplysia* neuronal CPEB were initially described in the context of the yeast model using a CPEB translation reporter as a read-out of activity [13]. When expressed in yeast, *Aplysia* CPEB maintains its function as a translational regulator and employs a prion mechanism to create stable, finely tuned, and self-perpetuating activity states controlling translation [13, 20]. While the prion-based activation of CPEB is evolutionarily conserved, little is known concerning the mechanism that induces CPEB structural switching.

In fungi, the chaperone machinery largely regulates protein folding and prion switching. Transient alteration in the chaperone machinery function can induce prion protein conformational switching, and this conformation persists for many generations [21-23]. The best-characterized regulator of prion switching in fungi is the Hsp104 chaperone. Transient perturbation of Hsp104 chaperone function either by the introduction of chemical or genetic Hsp104 inhibitors or by overexpression of the wild-type Hsp104 has been demonstrated to regulate the conformational switching of different prions and induce a heritable epigenetic trait [21-23]. Despite its critical role in protein folding homeostasis in fungi, Hsp104 has no known metazoan ortholog, suggesting that other chaperones have subsumed its function. For example, the chaperone Hsp70 also mediates some prion switching events in fungi [24-26] and is conserved in metazoans. Another highly abundant chaperone is Hsp90, which is evolutionarily conserved from bacteria to mammals. The role of Hsp90 in the cell extends far beyond the maintenance of protein homeostasis. For example, the ability of Hsp90 to bind a repertoire of metastable protein clients also enables Hsp90 to facilitate the evolution of diverse new biological phenotypes in different organisms [27-29]. Recent evidence suggests that the Hsp90 chaperone plays a role in prion switching as well [30-32].

In this work, using a proteomic approach, we reveal that the Hsp90 chaperone is a regulator of persistent CPEB translation activation. We then used a reporter of CPEB translational activation in yeast [13, 20] to elucidate an endogenous yeast mechanism through which transient Hsp90 suppression induces mRNA translation that persists for many generations. This work demonstrates that Hsp90 can regulate inheritance of a particular state of translational activity, elucidating an additional dimension to the role of Hsp90 as a major regulator of phenotypic diversity.

## Results

### Exploring the interactions of translation regulating proteins with the Hsp90 chaperone

We set out to explore how HSP90 regulates translation by characterizing the protein:protein interaction between HSP90 and a large library consisting of more than 1200 proteins that were shown to regulate translation or were associated with translation regulating processes based on Pfam nucleotide binding domains and previous experimental characterization [33, 34] (sup table 1). To quantify their interaction with Hsp90, we used LUMIER (luminescence-based mammalian interactome mapping) [35] (Fig 1A). In this assay, each protein from the library (bait) was Flag-tagged and overexpressed in HEK293T cells stably over-expressing HSP90 fused to Renilla luciferase. Following immunoprecipitation of the flag epitope, the protein-protein interaction was determined by the intensity of luminescence (Figure 1A). The distribution of interaction strength across all 1247 proteins (some with multiple isoforms) examined revealed that translation-related proteins bound HSP90 rather infrequently, less overall than previously described for transcription factors [35] (Figure 1B). We re-sequenced the plasmids encoding the proteins interacting with HSP90 and validated the Hsp90 interaction for these proteins. The HSP90 interacting proteins can be categorized as mRNA-binding, tRNA, rRNA and ribosomal biogenesis as well as other miscellaneous proteins (Figure 1C).

**Figure 1.**
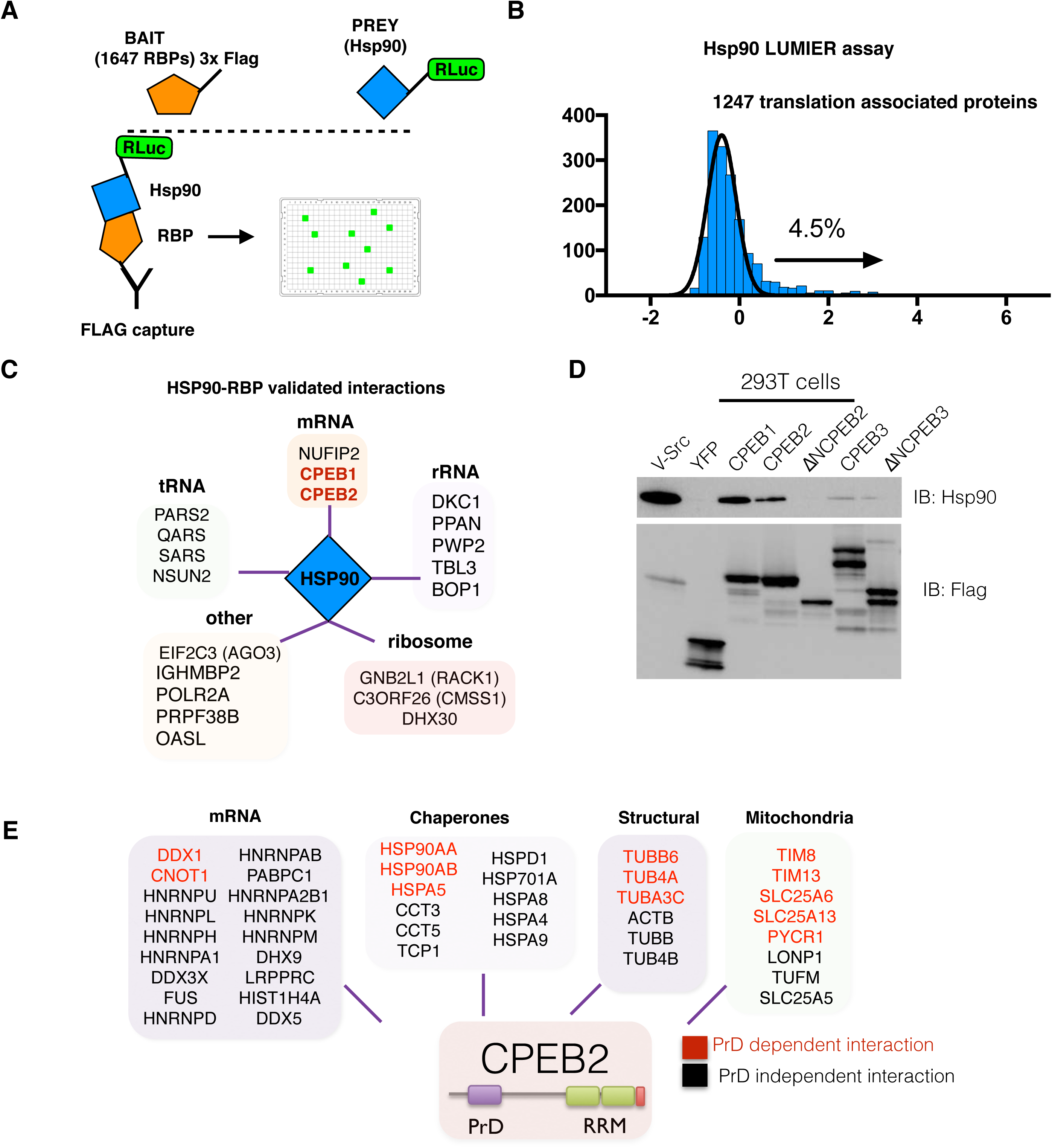
Proteomic approaches reveal a strong interaction of HSP90 with the CPE element binding protein 2 (CPEB2). (A) Schematic representation of the LUMIER assay for high throughput characterization of protein-protein interactions. 1647 Flag-tagged bait proteins (RBPs) are over expressed in 293T cells over expressing Hsp90 tagged with renila (Prey). Flag capture is performed in lates and the degree of protein-protein interaction is assessed by luminescence intensity. (B) Distribution of the Z-scores of all examined RBP interactions with Hsp90. 4.5% of the 1247 unique proteins (1647 with different isoforms) analyzed RBPs have a Z-score > 1.5. (C) Schematic of the classification of hits from the LUMIER assay that were validated to strongly bind HSP90 (raw data in Fig. S1). The CPE-element binding (CPEB) proteins are marked in red. (D) Flag-tagged CPEB1-3 proteins, CPEB2-3 lacking their PrD (DNCPEBs) and GFP and v-Src (used as controls) were over expressed in 293T cells and lysates immunoprecipitated with anti-flag beads. Interaction with the endogenous HSP90 was evaluated by immunoblotting with the indicated antibodies. (E) Schematic representation of proteins shown to associate with CPEB2 by mass spectrometry when over expressed in 293T cells. Proteins are classified according to the most abundant categories (Chaperones, Mitochondrial, Structural or mRNA-associated proteins). In red are the specific protein-protein interactions that were lost when the PrD domain of CPEB2 was deleted.

Interestingly, two CPE element binding proteins, CPEB1 and CPEB2, were among the strong Hsp90-binding proteins. We validated the interaction of both CPEB1 and CPEB2 with Hsp90 by performing a direct immunoprecipitation assay (Fig 1D, S1). The binding of CPEB2 to Hsp90 was dependent on its predicted prion domain as CPEB2 lacking the N-terminal region containing the prion domain (ΔNCPEB2) lost the ability to interact with HSP90. Although other CPEB family members have well-categorized interacting proteins, fewer binding partners have been described for human CPEB2. We therefore adopted a proteomic approach and examined the protein interaction landscape of CPEB2 in human cells. Full-length CPEB2 and CPEB2 lacking the PrD (ΔNCPEB2) were expressed in HEK293T cells and lysates immunoprecipitated with anti-flag. Co-precipitating interactors were assessed by mass spectrometry (Fig 1E Sup table 2). Four major functional categories of CPEB2 interacting proteins were revealed: chaperones, mitochondrial proteins, cell structure proteins and mRNA processing proteins (Fig 1E). Consistent with our immunoblot assay, the mass spectrometry results show that the Hsp90 interaction depends on the CPEB2 PrD (Fig 1E, PrD-dependent interactions in red). Indeed, the Hsp90-CPEB2 interaction shows the highest dependence on the PrD, but is not the only protein with this characteristic (Sup table 2). Thus, CPEB2 strongly binds Hsp90 in a PrD-dependent manner.

### Prion-like characteristics of the human CPEB2 protein

We determined whether human CPEB proteins had prion-like properties when expressed in yeast. The four human CPEB paralogs all harbor homologous RNA-recognition motifs (RRMs) but possess distinct N-terminal regions (Fig. 2A). Overexpression of CPEB1, CPEB2 and CPEB3 was toxic to yeast cells (Fig. 2B). Curiously, this was not due to the prion domain; indeed, removal of the CPEB2 PrD increased CPEB2 overexpression toxicity (Fig. 2C). However, this trend was reversed when the PrD was deleted from CPEB3 and the homologous deletion had little effect on the toxicity of CPEB4, suggesting unique functions and/or interactions for each PrD within each CPEB paralog (Fig. 2C).

**Figure 2.**
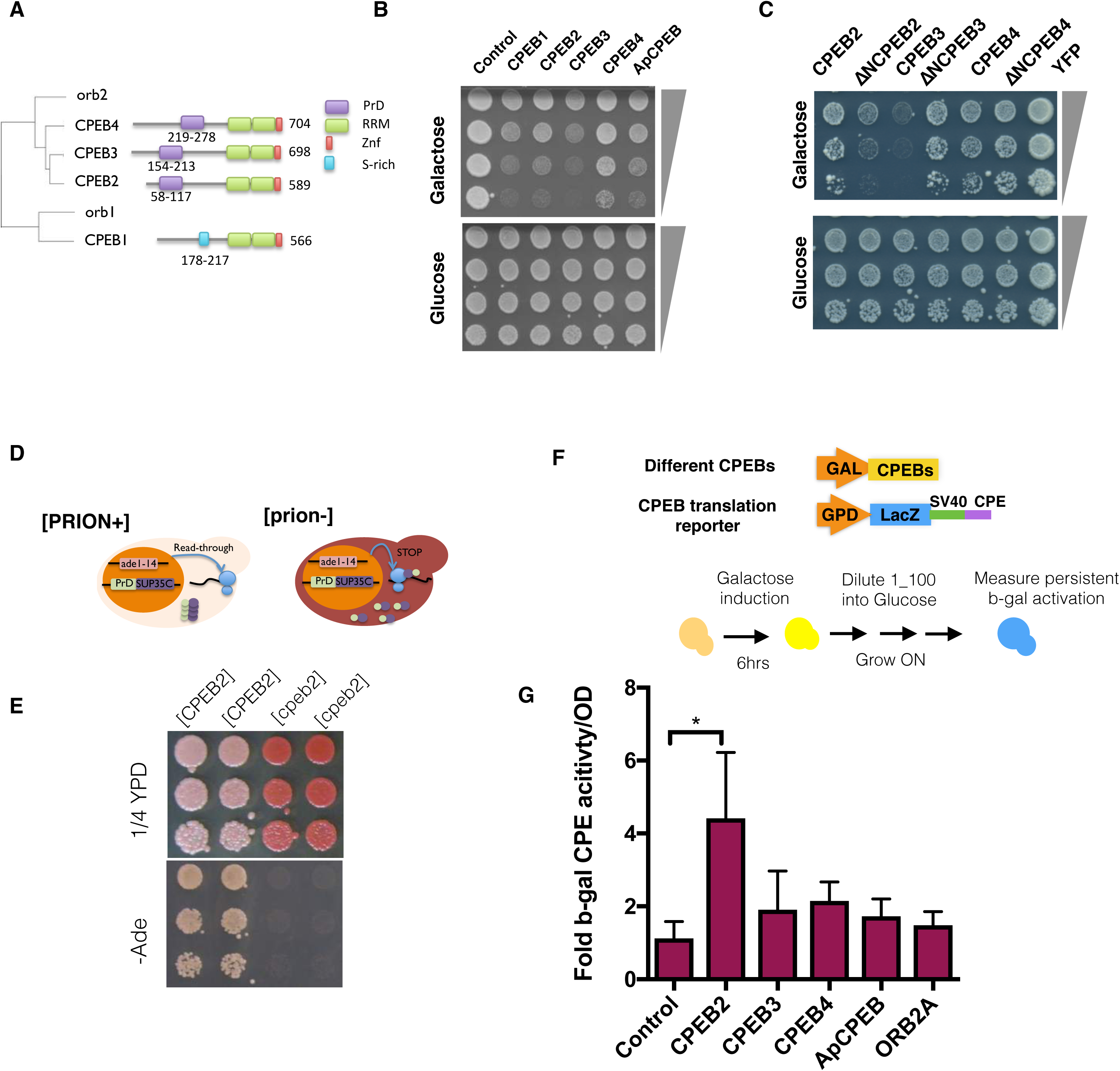
Characterizing human CPEB prion-like properties and Hsp90 interactions. (A) Phylogenetic tree representation of the human CPEBs, *Drosophila* (Orb2) and predicted yeast (Hrp1) orthologs. The functional domains (RRM-green, Zinc finger-red, Serine-rich-turquoise and PrD-purple) are shown on the four human CPEB homologs. (B-C) Yeast harboring Gal promoter full length (B) or PrD deleted (C) CPEBs tagged with mKate were grown overnight and then spotted onto expression-inducing plates (Galactose, upper) and uninducing plates (Glucose, lower) with 4-fold serial dilutions. (D) Schematic representation of the PrD-Sup35C assay. Cells in the [PRION+] state will read through the ade 1-14 mutation to produce a functional Ade product. This enables growth on plates lacking adenine. The resulting colonies are white due to lack of accumulation of red pigment. (E) Sup35C assay of the CPEB2 PrD. Shown is the ability of two distinct heritable conformations of the CPEB2PrD-Sup35C, white [CPEB2] and the red [cpeb2], to grow on medium lacking adenine and ¼ YPD. (F-G) Schematic representation of the experiment (F). 426 CPEBs under the control of the Gal promoter and tagged with mKate were introduced into the W303 alpha GEM strain. These yeast were then mated to a W303 strain expressing the CPEB β-gal translation reporter. The overexpression of CPEBs was achieved by incubating the cells ON in raffinose and then diluting them to an OD of 0.1 in 2% galactose. After 6 hours, the cells were diluted 1:100 into glucose-containing media and grown overnight (~ 7 generations). The OD600 and β-gal activities were then measured (G). The mean and SD of activity is presented, unpaired t-test was used for statistical analysis.

To examine the ability of the CPEB2 PrD to maintain a distinct heritable structural state, we exploited the modular nature of prion domains and the well-characterized [PSI^+^] phenotype in yeast. The prion domain of the Sup35 protein can be replaced by a prion domain of interest, producing a protein chimera that can form heritable phenotypes as does the wild type [PSI^+^]. The prion properties of such a chimera (PrD-Sup35C) can be visualized in yeast strains harboring an *ade1* nonsense allele. In the nonprion state these cells form red colonies and cannot grow on media lacking adenine. In the prion state, the chimeric protein aggregates and sequesters the Sup35 C-terminal translation termination activity, resulting in a read-through of the pre-mature stop codon of the ade1-14 allele, which leads to production of a functional Ade1 (nonsense suppression). In this [PRION+] state, the cells can grow in the absence of adenine and produce white colonies (Fig. 2D) [21].

To test the prion properties of the human CPEB2 protein, we overexpressed the CPEB2 PrD-SUP35C chimeric protein in cells lacking endogenous Sup35. In this isogenic background, the PrDCPEB2-Sup35C cells acquired two distinct, heritable states: exhibition of differential pigment accumulation when grown on rich media and growth on media lacking adenine (Fig. 2E, S2A). Furthermore, CPEB2 overexpressed in human cells exhibits SDS-resistant aggregation that is dependent on its prion domain (Fig. S2C). Thus, human CPEB2 exhibits many of the prion like characteristics previously described for the *Aplysia* CPEB [9, 13, 15, 20, 36]. The CPEB3 PrD exhibited prion properties similar to those of CPEB2 (Fig. S2A-B).

The hallmark of a prion phenotype is the ability to self-template and persist over many generations. In the context of prion-like proteins, their transient overexpression should be sufficient to induce a persistent phenotype [37]. The *Aplysia* neuronal CPEB, when expressed in yeast, can induce persistent activation of a translation reporter depending on its structural state [13]. The translation reporter has a cytoplasmic polyadenylation element (CPE) in its 3 UTR, which mediates increased β-gal (β-galactosidase) translation in response to CPEB binding [8, 15]. In the soluble state, CPEB is inactive, whereas in the prion aggregated form, CPEB induces activation of the translation reporter [13, 20]. We therefore examined the ability of the different CPEBs to induce persistent translation activation of the CPEB translation reporter upon transient overexpression in yeast. Expression of the CPEB proteins was induced for 6 hours and then the induction was switched off. The relative translation activation was measured after cells were grown for ~ 7 generations in the absence of CPEB production (Fig. 2F). Transient overexpression of human CPEB2 in yeast had the strongest persistent activation of the CPEB translation reporter (Fig. 2G).

### Transient Hsp90 inhibition induces persistent translation activation

The strong interaction of CPEB2 with Hsp90 prompted us to explore whether perturbation of Hsp90 activity could alter the prion-like CPEB-mediated translation regulation in yeast. We explored if transient Hsp90 inhibition would induce the prion-like properties of CPEB2 and promote a persistent increase in translation of the reporter even when the Hsp90 inhibitor is absent. Cells overexpressing CPEB2 were grown in the presence or absence of the Hsp90 inhibitor (radicicol) for 10 generations and then grown on media free of Hsp90 inhibitors. Transient inhibition of Hsp90 was sufficient to induce a persistent effect on CPEB2-mediated (β-gal translation when cells were grown in liquid media (Fig. 3A) and on solid media (Fig. 3B). Thus, suppression of Hsp90 activity augments the CPEB2-mediated translation activation in a self-sustaining manner.

**Figure 3.**
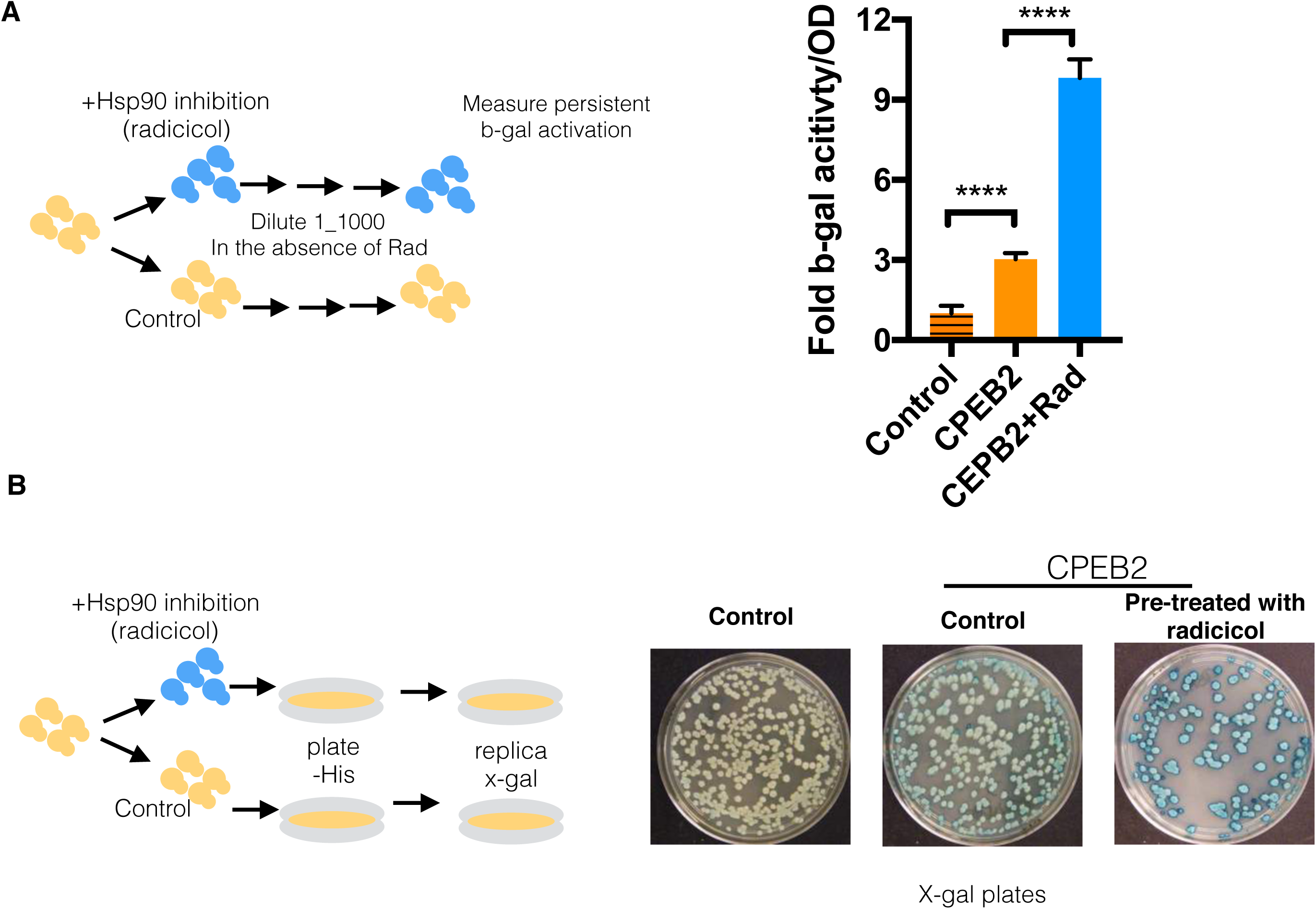
Transient suppression of Hsp90 induces a persistent translation activation. (A) Cells were grown overnight in media with or without 5uM radicicol. The cells were then diluted 1:1000 and grown for an additional 24 hours in media free of Hsp90 inhibitors to assess the persistent effect of transient Hsp90 inhibition on translation as measured by β-gal activity. The mean and SD of activity is presented, unpaired t-test was used for statistical analysis. **** p<0.0001 (B) CPEB2-overexpressing cells that harbor the CPEB translation reporter were treated with 10uM radicicol overnight and then plated on plates lacking radicicol. Cells were further replica plated on x-gal-containing plates to visualize the β-gal activity. Cells harboring the translation reporter but without CPEB2 overexpression were used as a control.

### Endogenous activation of the CPEB translation reporter in yeast in the absence of exogenous CPEB

Strikingly, transient Hsp90 inhibition had a significant effect on the persistent translation of the CPEB translation reporter even in the absence of exogenous CPEB (Fig. 4A). Although in the absence of exogenous CPEB the activation of the translation reporter was lower it still exhibited persistent high activity upon transient Hsp90 inhibition. This suggests the existence of an endogenous yeast mechanism that is affected by Hsp90 and regulated in a fashion similar to the regulation of exogenous CPEB. Suppression of Hsp90 activity for ten generations induced a persistent translation output in cells that was observable after growth on solid media without Hsp90 inhibitor for approximately 15 generations, as indicated after replica-plating onto x-gal plates (Fig 4B). Thus, even in the absence of exogenous CPEB, inhibition of Hsp90 induced a persistent translation output in cultures grown in either liquid or on plates.

**Figure 4.**
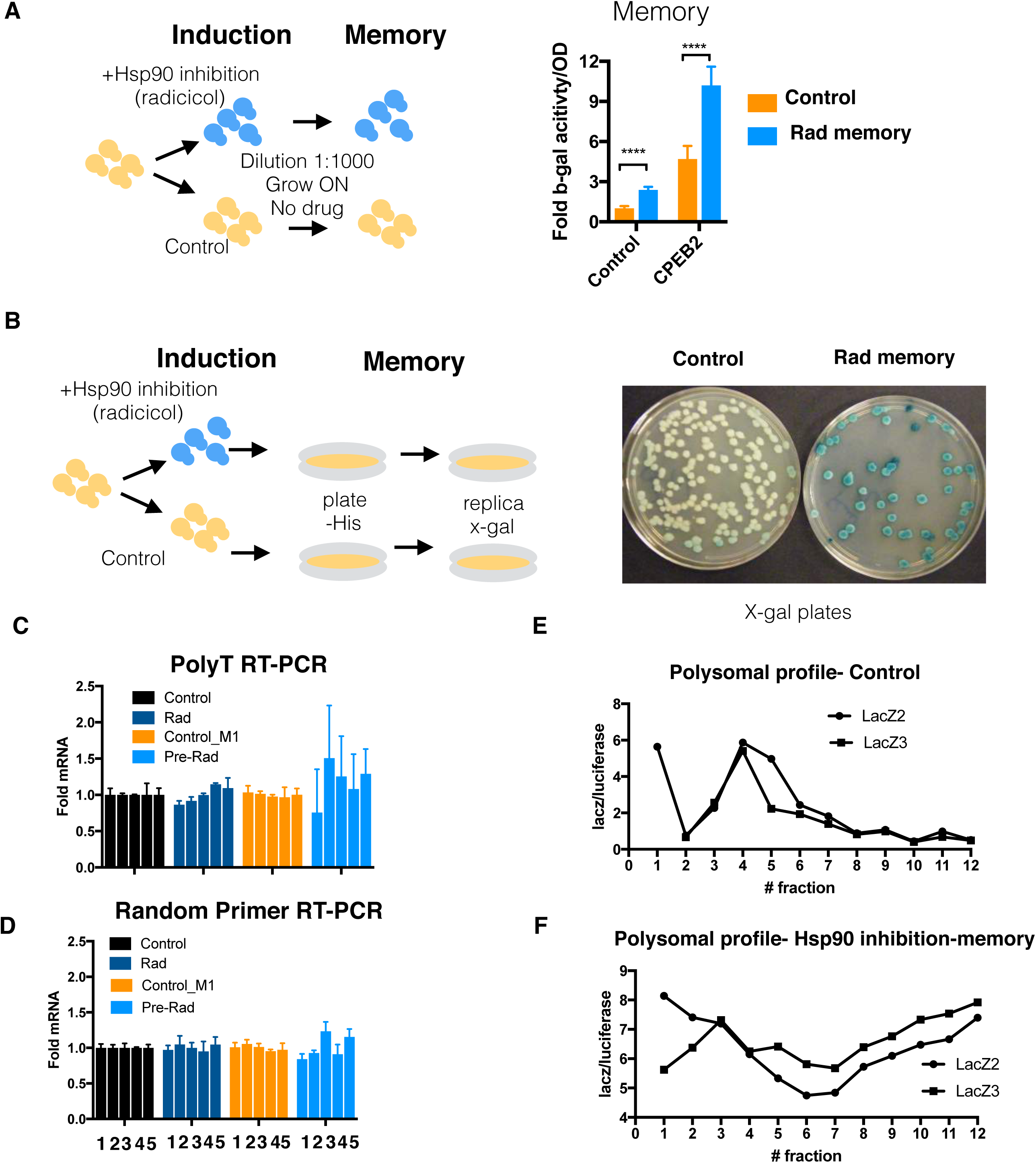
Transient suppression of Hsp90 activity induces a persistent translation activation in the absence of exogenous CPEB over expression. (A) Control and CPEB2-overexpressing cells were grown overnight with and without 5uM radicicol. Cells were then diluted 1:1000 and grown 24 hours in media lacking Hsp90 inhibitors The persistent effect of transient Hsp90 inhibition on translation was examined by measuring the β-gal activity. Results represent the mean values +- SD. Paired t-test analysis was conducted **** p< 0.001 (B) Cells were grown in liquid in the presence or absence of 5 uM radicicol and then plated on plates devoid of the Hsp90 inhibitor. After cells formed colonies, the plates were replica plated onto x-gal-containing plates to assess the persistence of β-gal activity. (C-D) The relative levels of β-gal open reading frame mRNA in the poly-A selected mRNA pool PolyT RT-PCR) (C) and total RNA pool (Random Primer RT-PCR) (D) in cells that were either treated with radicicol (rad) or control (Control) and then diluted to grow 10 generations in the absence of the drug (Pre-Rad) and (control-M1), respectively. (E-F) The distribution of β-gal mRNA across the polysomal fractions of control cells and cells that grew out from a 1:1000 dilution of a 10uM radiciciol-treated culture (Hsp90 inhibition-memory). LacZ2 and LacZ3 are two distinct primer sets used to amplify the β-gal mRNA and normalized to Luciferase (that was spiked into each fraction prior to analysis).

We next explored the effect on the mRNA levels of the CPEB translation reporter (β-gal) upon treatment with an Hsp90 inhibitor and the persistent effect of this exposure (“memory state”) compared to control. To do so, we initially performed quantitative PCR on the β-gal open-reading frame on both total RNA and the poly-A selected fraction. In both cases, no significant changes in mRNA levels were observed under any of the conditions (Fig 4C-D). We then performed polysomal fractionation followed by specific probing of the CPEB reporter (Figure 4E-F). In the memory state, β-gal mRNA shifted to the heavier polysomal fractions (Fig 4F), indicating greater translational throughput. Thus, the β-gal expression induced following transient Hsp90 inhibition is, indeed, due to increased translation with no detectable increase in the levels of β-gal mRNA.

In the process of these experiments, it came to our attention that the CPEB translation reporter used in the original report [13] included a predicted stem-loop structure in the 5’UTR that repressed its translation (LacZ-CPE). This structure was absent in the control reporter used in the same study (SYM-CAM) (Fig. S3A-F) [13]. To rule out the possibility that this stem loop might contribute to the CPEB-mediated translational activation, we made several adjustments to the original CPEB reporter. First, we removed the stem loop to eliminate the 5’UTR suppression. Second, we replaced the strong constitutive promoter with a weaker constitutive promoter (TDH3 promoter to SUP35 promoter) to decrease the level of transcription. Third, we replaced the long CPE-containing 3 UTR (Fig. S3A) with a short CPE element taken from the ADH1 3’UTR from *C.albicans*. Transient Hsp90 suppression was effective in inducing the persistent activation of β-gal in both the original and modified versions of the CPEB translation reporters (Fig. S3D-H). Thus, the induction of β-gal translation is not affected by altering promoters, or translation initiation inhibition in the form of a stem loop in the 5 UTR, suggesting that the regulation indeed arises from the 3 UTR.

### Translation activation induced by transient Hsp90 inhibition is reversible

In yeast, prion conformational switching is a rare event and has a low frequency of spontaneous reversion [38]. Reasoning that the activation of the CPEB translation reporter would also show trans-generational persistence, we tracked the persistence of reporter translation induction by transient Hsp90 inhibition to determine its reversibility. We treated cells with an Hsp90 inhibitor or DMSO as a control, grew out single colonies from both groups in the absence of the Hsp90 inhibitor and quantified the levels of the translation reporter for a span of almost 40 generations (schematic representation in Fig. 5A). Persistent activation of translation was significant after 20 generations and then gradually declined. Persistent activation remained significant for up to 30 generations, but not after 40 generations (Fig. 5B-C). Exploring the distribution of single colonies (Fig. 5C) further emphasized that the induction of translation and persistence were not uniform across clones despite all having been derived from a single clone prior to Hsp90 inhibition. The loss of translational memory after 40 generations indicates that this phenomenon is distinct from canonical prions as these would be expected to persist indefinitely in the lineage of most activated cells.

**Figure 5.**
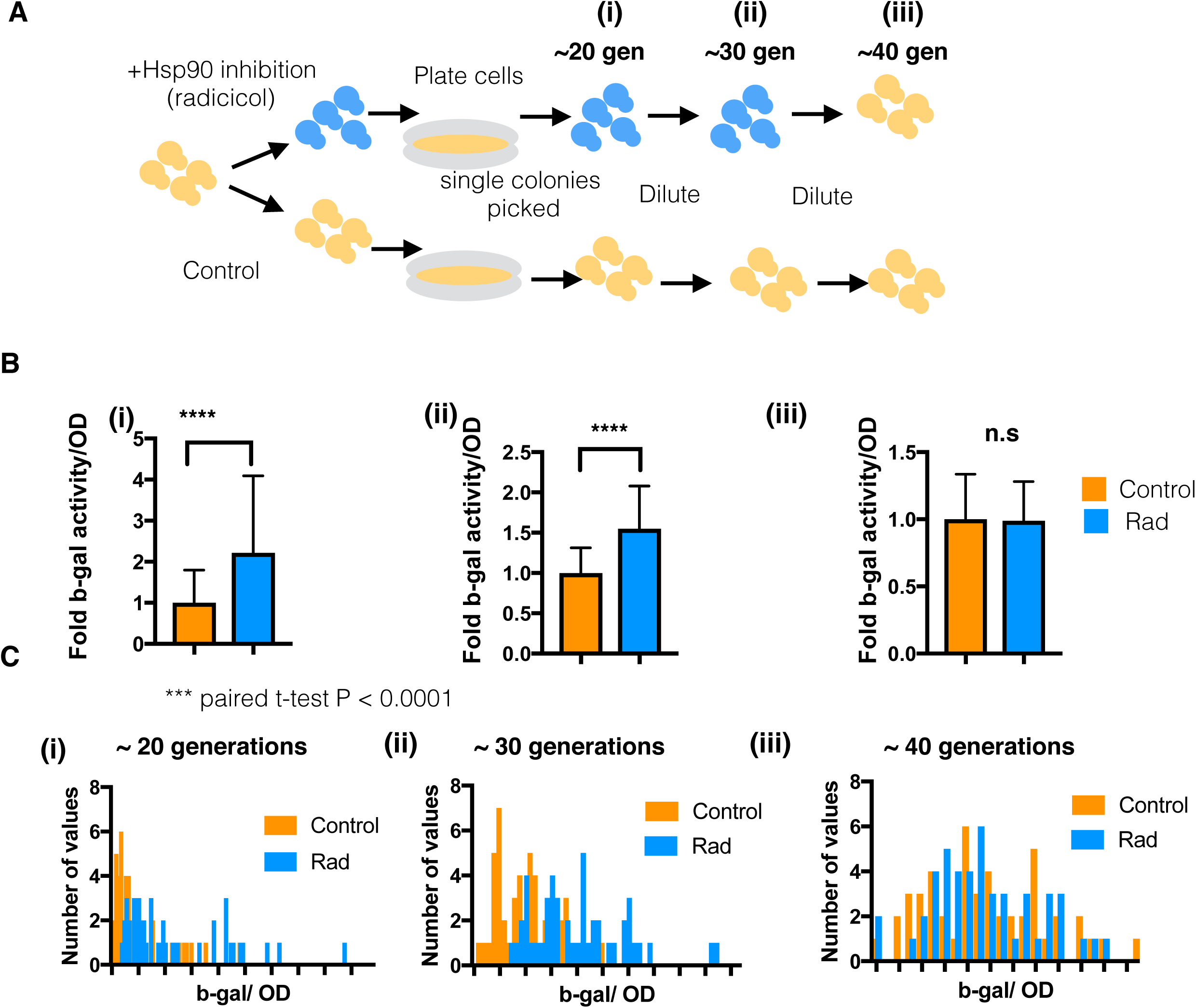
Endogenous activation of the CPEB translation reporter in yeast in the absence of exogenous CPEB. Cells were grown in the presence or absence of 10uM radicicol and then plated on plates devoid of the Hsp90 inhibitor. After cells formed colonies, the plates were replica plated onto x-gal-containing plates to assess the persistence of β-gal activity. (A-C) Cells were grown in the presence or absence of 10uM radicicol and then plated on plates devoid of the Hsp90 inhibitor. 48 individual colonies were picked and β-gal activity assessed after overnight growth (i). β-gal activity was further measured after dilution (1:1000) and an additional 24-hour growth period (ii), and again after sequential dilution (1:1000) and another 24-hour growth period (iii). Paired t-tests were conducted for statistical analysis. The schematic representation of the experiment (A), the median +-SD of the β-gal activity (B) and the overall frequency distribution of the individual colonies examined (C) are presented for the different time points. *** p<0.001.

### Aggregation of the 3’UTR-processing protein Hrp1 induces the CPEB translation reporter

Based on our finding that transient Hsp90 inhibition results in persistent activation of the CPEB translation reporter, we hypothesized that other modifications of protein homeostasis might elicit translational activation. Hrp1, a component of the 3 UTR-processing complex, was previously suggested to be the functional CPEB homolog in yeast [20]. Like CPEB, Hrp1 contains PrD and RRM domains and was recently shown to be highly aggregation prone [39]. To determine whether induced aggregation of Hrp1 promoted activation of the CPEB translation reporter, we assessed the effect of expressing an aggregation-prone mutant of Hrp1 (PY-mutant, [40]). Indeed, overexpression of the Hrp1 PY-mutant resulted in detectable aggregates (Fig. S4A). We further introduced the Hrp1 PY-mutant into the yTRAP (yeast transcriptional reporting of aggregating proteins) reporter system [39] (Fig. 6A). As expected, the expression of wt Hrp1 resulted in increased fluorescence compared to the PY-mutant (Fig. 6B), which in the yTRAP system indicates that the wt protein has greater solubility. This change in yTRAP signal was not due to decreased levels of the protein (Fig. S4B). Next, we introduced the CPEB translation reporter into the Hrp1 yTRAP strains to simultaneously monitor the Hrp1 aggregation state and the levels of translation. Under these conditions, the strains expressing wt Hrp1 exhibited a basal level of translation reporter activity, while the strains expressing the mutant aggregated form of Hrp1 exhibited heightened translational output as indicated by increased β-gal activity (Fig. 6C). These results suggest that the aggregation of a component in the 3’UTR processing machinery can mediate the induction of translation of the CPEB reporter.

**Figure 6.**
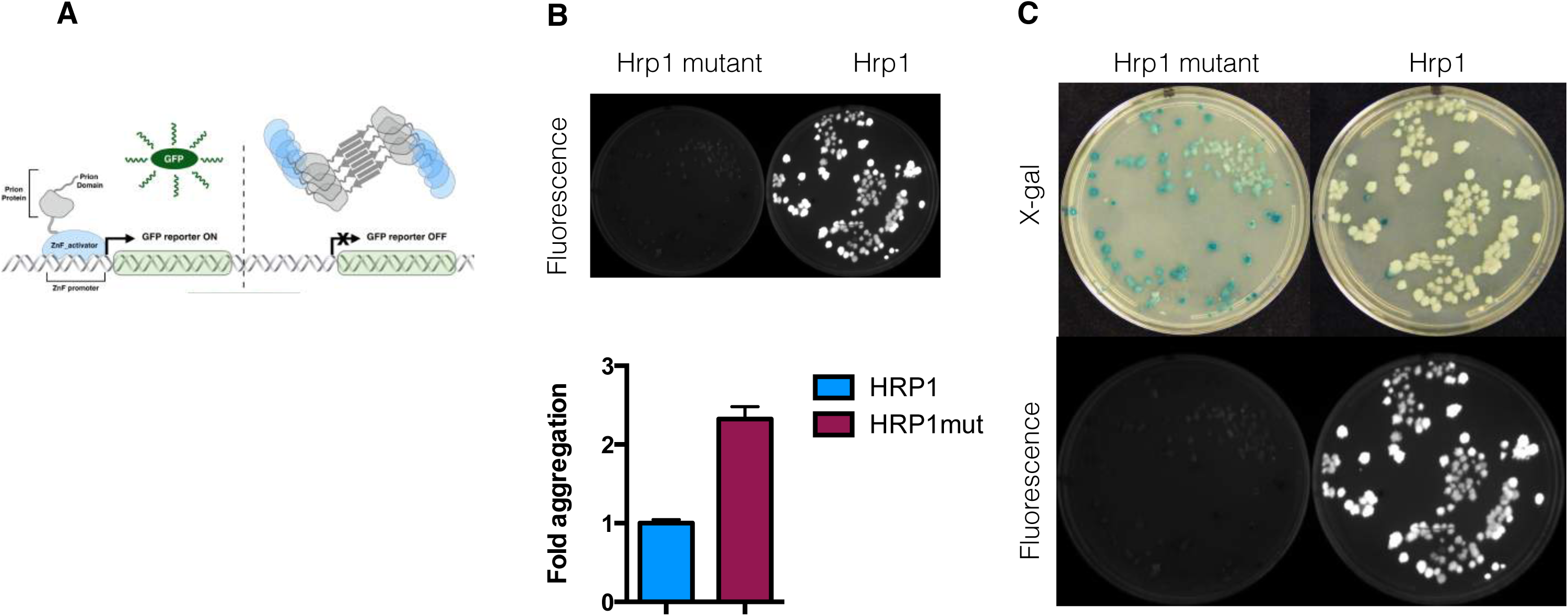
Aggregation of the 3’UTR-processing protein Hrp1 induces the CPEB translation reporter. (A) Schematic of yTRAP sensor mechanism (B) Visualization (top) and quantification (bottom) of fluorescence in cells harboring the wild-type (wt) and PY521/2AA mutant (mut) Hrp1 yTRAP sensor. (C) Cells harboring the wild-type or mutant Hrp1 yTRAP sensor and the CPEB translation reporter were replica plated onto x-gal plates and the fluorescent intensity and β-gal activity are shown.

### Perturbation of vacuole signaling induces the CPEB translation reporter but does not lead to translation memory

A more detailed time course analysis of translation reporter activation following Hsp90 inhibition revealed that activation peaked during the beginning of an alteration in growth rate associated with the yeast diauxic shift (Fig. 7A). Surprisingly, after removal of the Hsp90 inhibitor, this synchronous translation activation persisted, suggesting that both the original induction of translation and the persistent induction of translation depend on a specific cell state (Fig. 7A, bottom). One characteristic of the diauxic shift is that, during this time, the vacuole is the source of much of the nutrient signaling [41, 42]. To explore whether vacuolar signaling is involved in regulating the CPEB translation reporter, we genetically deleted various components of vacuolar pumps and measured the effects on translation output (Fig. 7B). Interestingly, genetic suppression of Vma1, a critical component of the V-ATPase complex, increased translation of the CPEB reporter (Fig. 7B). Hsp90 inhibition did not induce apparent Vma1 aggregation or mislocalization (Fig. S5A-C) but rather resulted in reduced protein levels of Vma1 (Fig. S5C). Deletion of other components of the V-ATPase complex also induced the translation reporter, suggesting that the effect on translation is not specifically due to regulation of Vma1 but rather due to impairment in V-ATPAse functionality (Fig. S5D). Further examination of organelle morphology following Hsp90 inhibition revealed some mitochondrial structure abnormalities (Fig. S5E), suggesting that translation activation induced by Hsp90 inhibition might involve the vacuolar-mitochondrial axis.

**Figure 7.**
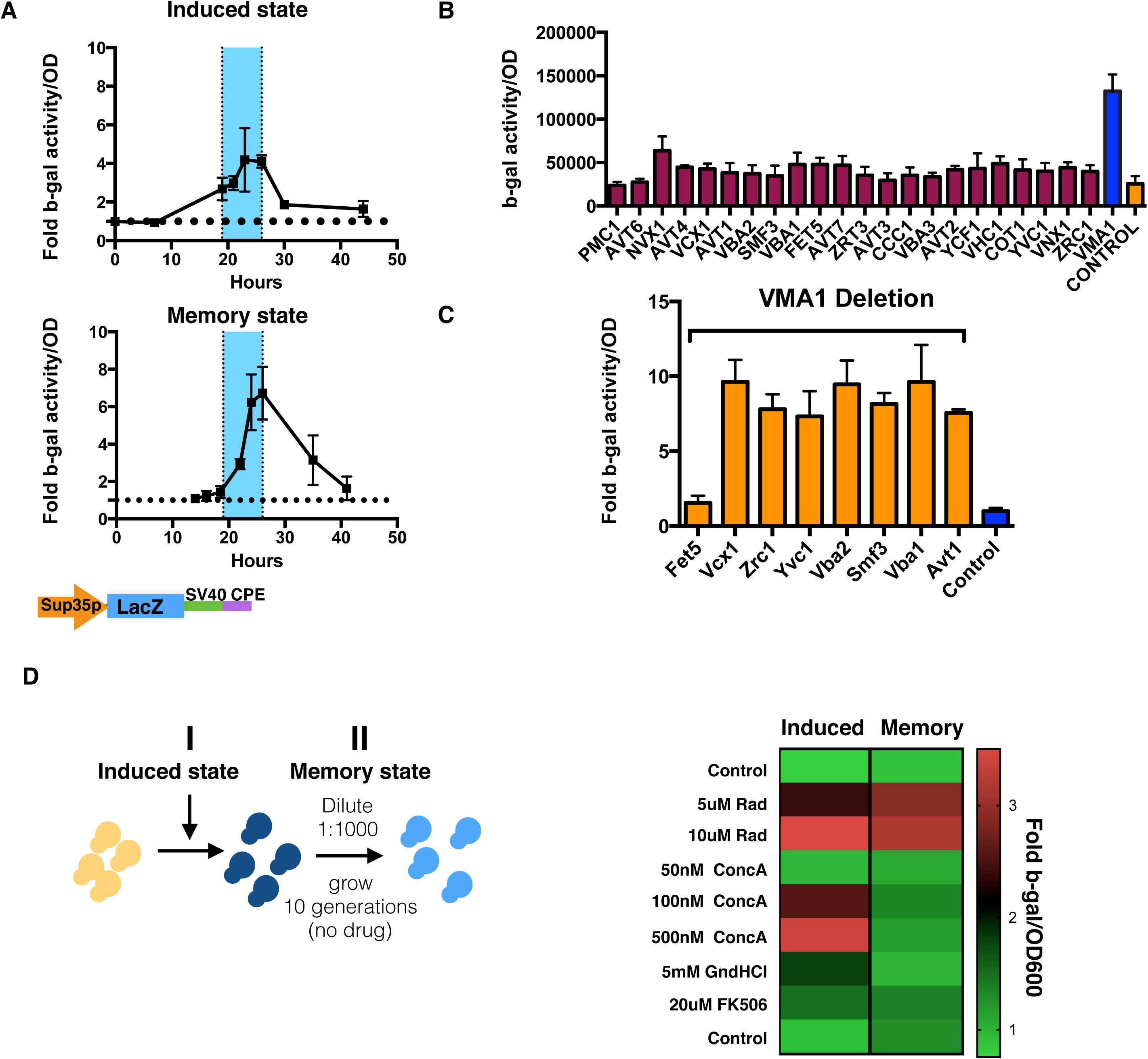
Perturbation of the vacuole-mitochondria axis induces the activation of translation but the effect is not persistent. (A) The time course of fold activation of β-gal was analyzed when cells were grown in the presence or absence of the Hsp90 inhibitor radicicol (upper, induced state) and after cells were diluted 1:1000 and grown in the absence of the drug (lower memory state). (B) β-gal activity was analyzed in strains deleted for different vacuole-associated proteins that harbored the CPEB translation reporter. (C) The indicated vacuolar pumps were overexpressed on the background of VMA1 deletion and the activity of the CPEB translation reporter was analyzed. (D) The induction and persistence of β-gal activity was analyzed in cells harboring the CPEB translation reporter. Cells were treated with the indicated concentration of drugs (ConcA-concanamycin A (V-ATPase inhibitor), GndHCl (Hsp104 inhibitor), FK506 (Calcineurin inhibitor). The persistent effect was analyzed after cells were diluted 1:500 and grown in the absence of the drugs for 24 hours.A model of activation of the yeast [CPEB] prion translation phenotype

The V-ATPase function of acidifying the vacuole can affect a large subset of vacuole functions. For instance, many vacuolar pumps depend on the proton gradient generated by the V-ATPase complex. To identify signaling components downstream of Vma1, we over-expressed specific vacuolar pumps in the background of the Vma1 deletion to determine if any would reverse the β-gal activation observed in that strain. Overexpression of the vacuolar iron/copper transporter Fet5 completely eliminated the Vma1 deletion-induced β-gal activation (Fig. 7C). Fet5 is a multi-copper oxidase that plays a role in iron transport [43] and recently was shown to regulate iron recycling during diauxic shift [44]. Culturing cells in media depleted of iron and copper was sufficient to increase the expression of β-gal from the CPEB translation reporter and the effect was additive to radicicol activation (Fig. S5F). Thus, signals associated with iron homeostasis that arise from the vacuole and are potentially mediated by Fet5 affect the translation of the β-gal CPE-reporter.

If Hsp90 inhibition induces the persistent translation of the CPEB reporter solely through Vma1 suppression, we would expect a similar outcome with transient suppression of the V-ATPase complex. Indeed, the V-ATPase inhibitor Concanamycin A (ConcA), like the Hsp90 inhibitor, induced activation of the translation reporter (Fig. 7D). However, persistent translation activation was only achieved with Hsp90 suppression (Fig. 7D). Thus, Hsp90 inhibition is specific and unique in inducing and promoting a heritable change in protein translation.

## Discussion

In this work, we establish an intriguing phenomenon of Hsp90-regulated, non-genetically inherited translation memory. We first established a strong link between the human CPE element binding protein 2 (CPEB2) translation regulation and the Hsp90 chaperone by surveying the protein-protein interactions of HSP90 with a large library of RNA-binding proteins. We discovered a strong protein-protein interaction between CPEB2 and HSP90 that is mediated by the prion-like domain of CPEB2. The functionality of this protein-protein interaction is revealed upon Hsp90 inhibition: even transient suppression of Hsp90 is sufficient to induce the activation of the CPEB translation reporter in the yeast model. This translation activation is persistent and gradually declines over 40 generations. Moreover, transient suppression of Hsp90 induces the persistent translation activation of the CPEB translation reporter even in the absence of exogenous CPEB overexpression, suggesting yeast possess an endogenous CPEB-like molecule that mediates translational memory. Although other perturbations, including iron-copper depletion and V-ATPase deletions, can affect the translational reporter, Hsp90 inhibition was the only identified stimulus that triggered persistent activation over many generations of growth. Thus, another Hsp90 interactor likely acts like CPEB2 to induce the translation of mRNAs.

The evolutionarily conserved family of CPEB proteins can be subdivided by the presence or absence of a prion domain. In *Aplysia, Drosophila* and mice, the prion domain has distinct functions in promoting CPEB activity [9, 13-18, 36]. In humans, there are four homologs of CPEB; three (CPEB2-4) contain a predicted prion-like domain and one does not (CPEB1). Using the translation assay originally designed to characterize the prion properties of the *Aplysia* CPEB [13], we show that the expression of human CPEB2 yields the strongest heritable activation of translation. The prion domains of both CPEB2 and CPEB3 enable prion-like inheritance. However, the function of these domains seems to be specific and unique in the context of the whole protein as its absence increases the toxicity of CPEB2 but reduces the toxicity of CPEB3. Moreover, our proteomic analysis revealed that the PrD of CPEB2 is crucial for mediating specific protein-protein interactions, including interactions with the chaperone Hsp90, proteins involved in mRNA translation regulation, tubulin and mitochondrial translocation proteins (TIM8 and TIM13). Interestingly, CPEB3 protein-protein interaction fall largely within the same functional interaction as for CPEB2 however different PrD-dependent protein-protein interaction was observed for CPEB3.

The strong interaction between the PrD of CPEB and Hsp90, along with the profound induction of the translational memory upon Hsp90 inhibition suggests that Hsp90 is a key mediator of the prion-like phenotypes of CPEB. CPEB2 overexpression was sufficient to promote the activation of translation as previously shown for the *Aplysia* CPEB [13]. However, the Hsp90 inhibition further enhanced this activation of translation in a heritable fashion. The surprising finding is that suppression of Hsp90, even in the absence of exogenous CPEB, can induce a persistent and inherited translation phenotype in yeast. Although this activation is lower in magnitude than the one observed in the presence of CPEB, it is nevertheless sufficient to promote a heritable translation phenotype that declines gradually over 40 generations.

We believe these data are best explained by the existence of an endogenous yeast RNA-binding protein with prion-like properties with similar structure–function characteristics as described for the CPEB proteins [16-20, 45], and for simplicity will be termed [CPEB] prion. The enhancement of the [CPEB] phenotype by overexpression of exogenous CPEB and induced Hrp1 aggregation support a model of altered 3’UTR regulation. Although the aggregated Hrp1 induced the translation, we have no additional evidence to support the fact that Hrp1 is the yeast [CPEB] prion. However, the aggregation of an element from the 3’UTR processing complex could potentially induce the sequestration or co-aggregation of the actual [CPEB] prion as reflected by the induction of translation by aggregated Hrp1. Many of the yeast RNA-binding proteins contain segments resembling prion-domains and have the ability to form stable aggregates [21, 46-48]. In some cases, as shown for Whi3 and Rim4, aggregation is an integral part of their biological function [45, 49, 50]. Supporting the possibility that the [CPEB] prion phenotype is mediated by a protein-based mechanism mediated by an RNA-binding protein.

In our model, upon transient Hsp90 suppression, the endogenous yeast prion transitions from the [cpeb] translation inactive state to the [CPEB] prion state that self-templates additional functional aggregates resulting in persistent translation activation of specific mRNAs. This prion-like phenomenon replicates enough to provide 30 generations of phenotypic consequences without the exponential dilution that would occur for non-prion assemblies. However, ultimately this prion-like state is not stable and through either its own instability or the action of Hsp90, is eliminated after additional generations. The [CPEB] prion activity following Hsp90 inhibition peaks at the diauxic shift and is also strongly affected by the V-ATPase and iron-copper signaling. This might suggest that the induction of the [CPEB] prion protein expression or stabilization (or post-translational modification) is achieved at the diauxic shift stage and is regulated by lysosomal signaling. Perturbing this regulation alters the activation of the CPEB translation reporter but does not result in heritable [CPEB] prion state with persistent translation activity as observed in the case of Hsp90 suppression (Fig. 8). Thus, Hsp90 inhibition is unique in promoting the [CPEB] prion persistent translation activation revealing yet another mechanism through which this powerful chaperone enhances phenotypic variation.

**Figure 8.**
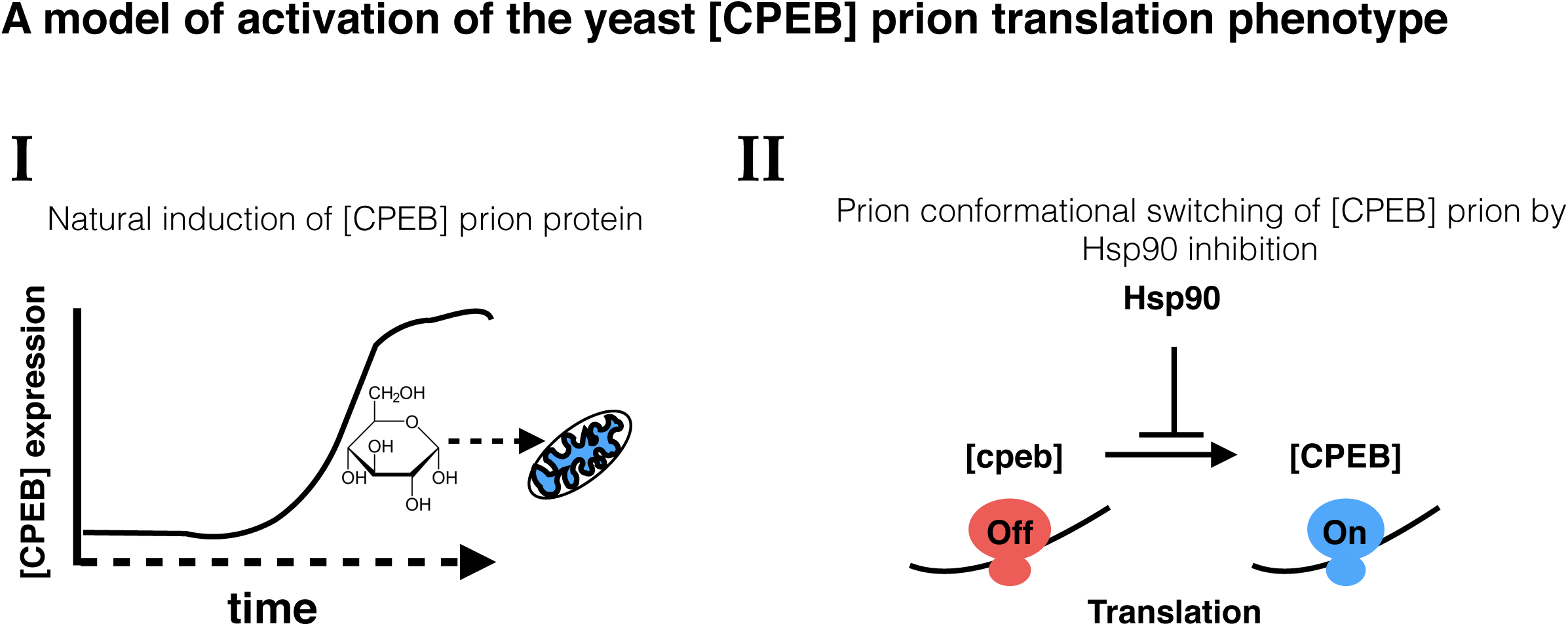
A model of Hsp90 inhibition-induced translation memory. (i) The protein mediating the expression of the [CPEB] prion phenotype is induced during the transition from glycolysis to mitochondrial-dependent metabolism and is affected by signaling from the vacuole. (ii) Suppression of Hsp90 induces the [CPEB] prion conformational switching. Increased prion switching with Hsp90 inhibition will enhance translation and enable epigenetic inheritance of an altered translation state. In this model in order for the [CPEB] prion translation phenotype to be induced there is a need for the activation of the mediator prion protein (i) and the Hsp90 induced switch to enable the persistence and inheritance of the [CPEB] prion translation state (ii).

## Acknowledgments

This work is dedicated to the special times and people of the Lindquist lab and to Susan Lindquist that enabled us to flourish in this amazing environment that tragically came to an end. Thanks to all the Lindquist lab members that stuck around the hard times and helped in many ways beyond the work itself, and the Lindquist lab alumni that reached out and gave much support. Special thanks to Linda Clayton for the constructive reading and comments and the general support over all the years. We would like to thank Gerald Fink for his supervision and constructive comments, Dirk Landgraf for the endogenously tagged fluorescent yeast strains and Kausik Si for the human yeast codon optimized CPEB PrD plasmids. P. Tsvetkov was supported by EMBO Fellowship ALTF 739-2011 and by the Charles A. King Trust Postdoctoral Fellowship Program. This work was supported by the Mathers foundation. S.L. was an investigator of the Howard Hughes Medical Institute.

## Methods

**Clones and Human cell lines** – To create the translation related gene library, genes with known RNA binding domains (pfam), or those annotated to have translation regulating function by Gene Ontology, Uniprot, and literature curated. The specific clones were collected from the human ORFeome [51], and those absent were cloned from cDNA by PCR. Clones were transferred into pcDNA3.1-based mammalian expression vector containing a C-terminal 3 × FLAG tag [35, 52] using Gateway recombination. Inserts were verified using restriction digestion (BsrGI, NEB) and Sanger sequencing. HEK293T cells Reporter cell lines stably expressing Renilla luciferase (Rluc)-tagged HSP90 (HSP90β) were previously described [35].

**LUMIER assay** − The LUMIER assay was conducted as previously described [35]. 3 × FLAG-tagged bait proteins were transfected into stable 293T cell line in 96-well format with polyethylemine (PEI). Two days after transfection, cells were washed in 1 × PBS and lysed in ice-cold HENG buffer (50 mM HEPES-KOH [pH 7.9], 150 mM NaCl, 20 mM Na_2_MoO_4_, 2mM EDTA, 5% glycerol, 0.5% Triton X-100). The lysate was transferred to 384-well plates coated with anti-FLAG M2 antibody (Sigma-Aldrich). Plates were incubated in cold room for 3 hr, after which plates were washed with the lysis buffer using an automated plate washer. Luminescence in each well was measured with an Envision plate reader using Gaussia FLEX luciferase kit (New England Biolabs). The normalized luminescence Z scores was used as a quantitative interaction measure.

**Immunoprecipitation** – HEK293T cells were transfected with 3xflag tagged bait proteins. 48 hours after transfection cells were washed in 1 × PBS and lysed in ice-cold HENG buffer. Proteins were immunoprecipitated using flag conjugated beads at 4°C for 3 hours. Beads were then washed in HENG buffer to remove non-specific interactions and immunoprecipitated proteins were removed from beads by addition of loading buffer and heating 5min 95°C. Samples were then loaded on Bis-Tris 4%–12% PAGE, and analyzed by western blotting on nitrocellulose membranes.

**SDD-AGE- SDD-AGE** was performed as detailed in [21]. A final concentration of 1% SDS was used in the loading buffer.

**Read through assay**- *ade1-14* read-through was assessed as detailed in [21]. Briefly, the CPEBPrD-Sup35c chimeras were individually expressed in the YRS100 strain (a derivative of YMJ584), where the endogenous Sup35 protein has been deleted, but the activity of this essential protein is covered by 416GPDSup35C. Then a plasmid shuffle was performed by plating the cells in 5-fluoroorotic acid (5-FOA). Therefore, the CPEB PrD-Sup35C was the only source of Sup35C activity in these cells.

**Polysome profiling-** Yeast lysates were prepared as described [53] with minor modifications. Briefly, samples were rapidly collected using filtration and ground to a fine powder using a Freezer/Mill (SPEX SamplePrep) and stored at −80 °C. Lysates were thawed and resuspended in lysis buffer (10mM Tris-HCl, pH 7.4, 5mM magnesium chloride, 1% Triton X-100, 1% sodium deoxycholate, 100mM KCl, 0.02U/μL Superase-IN, 2mM DTT, and 100μg/mL cycloheximide), and homogenized on a Disruptor Genie (Scientific Industries) for 1 minute. The supernatant was loaded onto a 10-50% w/v linear sucrose gradient containing 20mM HEPES-KOH, pH 7.4, 5mM MgCl2, 100mM KCl, 2mM DTT, 100μg/mL cycloheximide, and 0.02U/μL Superase-IN. Gradients were centrifuged for 2hr at 36,000 r.p.m. and 4°C and then fractionated by upward displacement using a Biocomp Gradient Station with continuous absorbance monitoring at 254nm. Luciferase RNA (Promega) for qPCR normalization was added to fractions of equal volume immediately after collection. Total RNA was extracted using TRI reagent (Ambion) according to the manufacturer’s instructions.

**Mass spectrometry analysis**-, immunoprecipitation was conducted as described above and analysis of proteins was conducted from the SDS-PAGE gel lanes for each sample were subdivided into 15 molecular weight regions. These gel bands were reduced, alkylated and digested with trypsin at 37°C overnight. The resulting peptides were extracted, concentrated and injected onto a Waters HPLC equipped with a self-packed Jupiter 3 μm C18 analytical column (0.075 mm by 10 cm, Phenomenex). Peptides were eluted using standard reverse-phase gradients. The effluent from the column was analyzed using a Thermo LTQ linear ion trap mass spectrometer (nanospray configuration) operated in a data dependent manner. The resulting fragmentation spectra were correlated against the known database using SEQUEST. Scaffold Q+S (Proteome Software) was used to provide consensus reports for the identified proteins.

**Creating the new CPEB translation reporters**- The original 315 LacZ CPE reporter was the original reporter described (ref), and the control SYM-CAM was also previously described (ref).

*Removing the loop from original LacZ-CPE reporter-* Removing the loop at the 5’UTR of the original LacZ-CPE reporter was done by digesting the plasmid with NcoI and SpeI, gel purification of the digested fragment and ligation. The plasmid was validated by sequencing to ensure that the plasmid retained the ORF sequence. This resulted in the LacZ-CPE no loop CPEB translation reporter.

*Changing the Leu2 cassette to HIS3-* The Leu promoter driving Leu2 gene was cut out from original reporter with BrgI and KapI digest. TEF promoter driving HIS3 gene with a TEF terminator were inserted by Gibson assembly following PCR reaction with the following primers.

His Fw Primer: gtttggccgagcggtctaagGACATGGAGGCCCAGAATAC

His Rv Primer: gaatttcatttataaagtttatCAGTATAGCGACCAGCATTC

*Replacing TDH3 promoter with loop with Sup35 promoter-* Sup35 promoter was used to replace the TDH3 promoter and the strong stem loop at the 5’ UTR of the 315 LacZ CPE reporter. This was done by PCR of the SUP35p using the following primers:

Fw Sup35p: ctcgccatttcaaagaatacCAACCACACAAAAATCATACAAC

Rv Sup35p: tcttttttggctccatggcaTGTTGCTAGTGGGCAGATATAG

The reporter was digested with HindIII and SnaBI and the Sup35 promoter was inserted using Gibson assembly. Resulting in the Sup35p Lacz-Sv40-CPE translation reporter. *Replacing the SV40 CPE 3’UTR with an ADH1 terminator -* initially a NheI digest site was introduced after the stop codons of LacZ to enable the manipulation of the 3’UTR

sequence. The NheI digest site was introduced by PCR of the backbone plasmid with the following primers:

Fw Backbone (NheI) (51-mer):

TGGTCTGGTGTCAAAAATAATAATAAgctagcCCGGGCAGGCCATGTCTGC Rv backbone (54-mer):

CTCTTCTTTTTTGGCTCCATGGCAgtttaaacTGTTGCTAGTGGGCAGATATAG The LacZ ORF was amplified with the following primers:

Fw LacZ (PmeI) (30-mer): gtttaaacTGCCATGGAGCCAAAAAAGAAG Rv PCR lacZ (34-mer): GCTagcTTaTTaTTATTTTTGACACCAGACCAAC The two amplified segment were assembled using Gibson assembly.

The Sup35p LacZ-SV40-CPE CPEB translation reporter with the new NheI digest site was digested with NheI SacII to remove the SV40-CPE 3’UTR and an ADH1 3’UTr was Amplified using the following primers:

Fw ADH1: gtctggtgtcaaaaataataataagctagcTAAGCAAATAGCTAAATTATATACG RvADH1:

cgaattggagctccaccgcggtggcggccgCAACTGTATAAGATAGTAATAAAAATATCG

The fragments were assembled by Gibson assembly method to yield the Sup35p-LacZ-ADH1 CPEB translation reporter.

**Vma1-mCherry tagging**- The tagging was performed as previously described [54] using pFA6a_mCherry_HPHMX as template and the following primers :

VMA1Cterminal tagging f1: CGGCCTTGTCTGATAGTGATAAGATTACTTTGGATG

VMA1Cterminal tagging r1: ATCAACCTGTAGGGTTCTATCGGTAGATTCAGCAAATCTTTCTTGCATAGTGCTCAAC

VMA1Cterminal tagging f2:

GTTTAAACGAGCTCGAATTCGATATATGTAGCATTTATCTTCTGGTATATTTGTTAG GTTTAAACGAGCTCGAATTCGATATATGTAGCATTTATCTTCTGGTATATTTGTTAG

VMA1Cterminal tagging r2: CTACCTCATAATGGATCTAAATTGCATACTAAT CTCAC

**Quantitative PCR analysis**- total RNA was purified via phenol/chloroform separation using phase lock tubes (five prime) followed by ethanol precipitation as previously described [55]. RT-PCR PCR was conducted with either poly-T primers or random primers using Superscript III (Thermo fisher) according to manufactures protocol. qPCR was performed with the primers for LacZ (set of 5 pairs) and the genes for normalization [56] (ALG9, TAF10, TFC1)

fLacZ1: ATC TTC CTG AGG CCG ATA CT

rLacZ1: CGG ATT GAC CGT AAT GGG ATA G

fLacZ2: CCA ACG TGA CCT ATC CCA TTA C

rLacZ2: TTC CTG TAG CCA GCT TTC ATC

fLacZ3: GTT GGA GTG ACG GCA GTT AT

rLacZ3: GCT GAT TTG TGT AGT CGG TTT ATG

fLacZ4: GCC GAA ATC CCG AAT CTC TAT C

rLacZ4: AGC AGC AGC AGA CCA TTT

fLacZ5: CAT GTT GCC ACT CGC TTT AAT

rLacZ5: GAA ACT GTT ACC CGT AGG TAG TC

ALG9

Fw: CACGGATAGTGGCTTTGGTGAACAATTAC

Rv: TATGATTATCTGGCAGCAGGAAAGAACTTGGG

TAF10

Fw: ATATTCCAGGATCAGGTCTTCCGTAGC

Rv: GTAGTCTTCTCATTCTGTTGATGTTGTTGTTG

TFC1

Fw: GCTGGCACTCATATCTTATCGTTTCACAATGG

Rv: GAACCTGCTGTCAATACCGCCTGGAG

**Sup figure 1.**
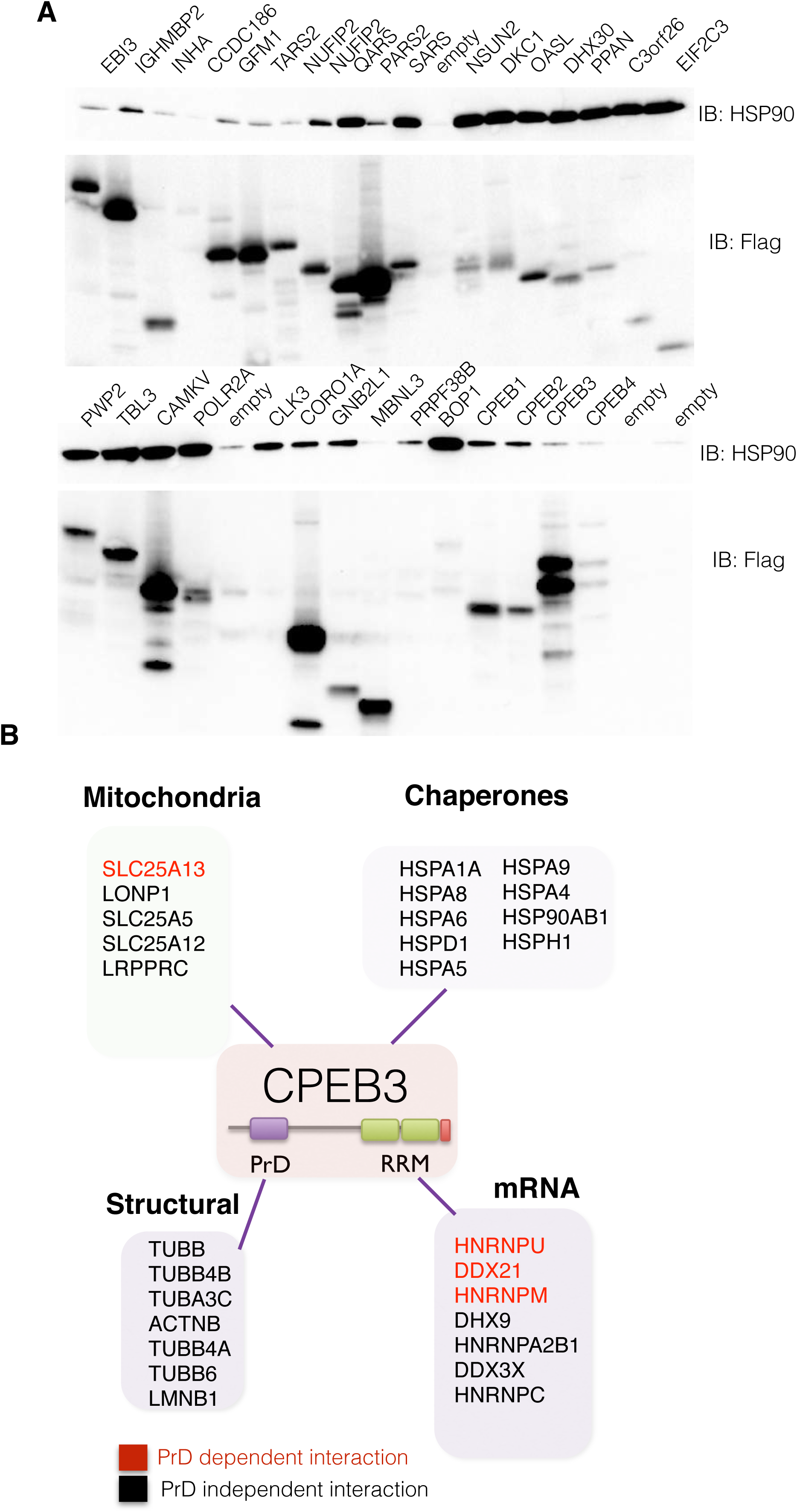
(A) The indicated flag-tagged proteins were over expressed in 293T cells and their binding to the endogenous Hsp90 was assessed following immunoprécipitation with anti-flag antibody and immunoblotting with anti-HSP90. (B) Schematic representation of proteins that were found to associate with CPEB3 when overexpressed in 293T cells. CPEB3-associating proteins are classified to the most prevalent categories (Chaperones, Mitochondrial, Structural or mRNA associated proteins). In red are the specific protein-protein interactions that were lost when the PrD of CPEB3 was deleted.

**Sup figure 2.**
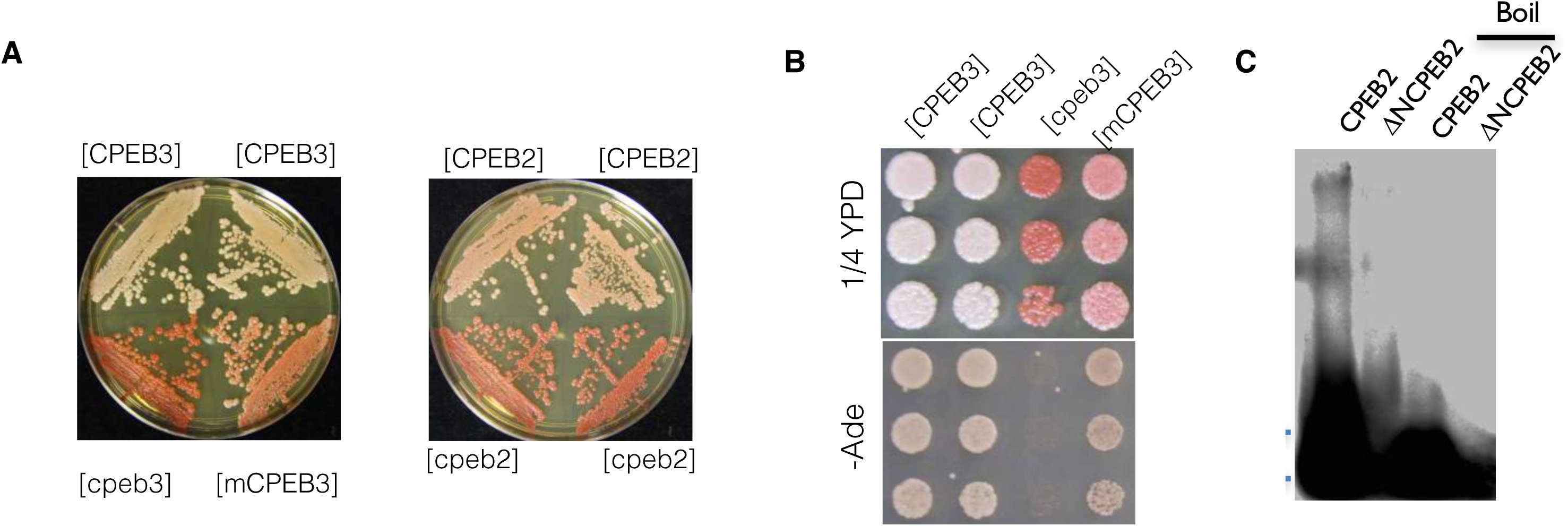
(A) Streak out of white [CPEB2/3] and red [cpeb2/3] PrD-SUP35C colonies reveals that the epigenetic state of each strain is maintained over many generations. CPEB3 exhibited an intermediate state [mCPEB3], with altered pigmentation but lack of ability to grow on -Ade. (B) Sup35C assay of the CPEB3 PrD. Shown is the ability of two distinct heritable conformations of the CPEB3PrD-Sup35C, white [CPEB3] and red [cpeb3], to grow on medium lacking adenine or on % YPD. (C) Cell lysates from 293T cells over expressing flag-tagged full length CPEB2 or the PrD-deleted CPEB2 were subjected to SDD-AGE analysis with and without boiling prior to loading. Proteins were detected by immunoblotting with anti-flag antibody.

**Sup figure 3.**
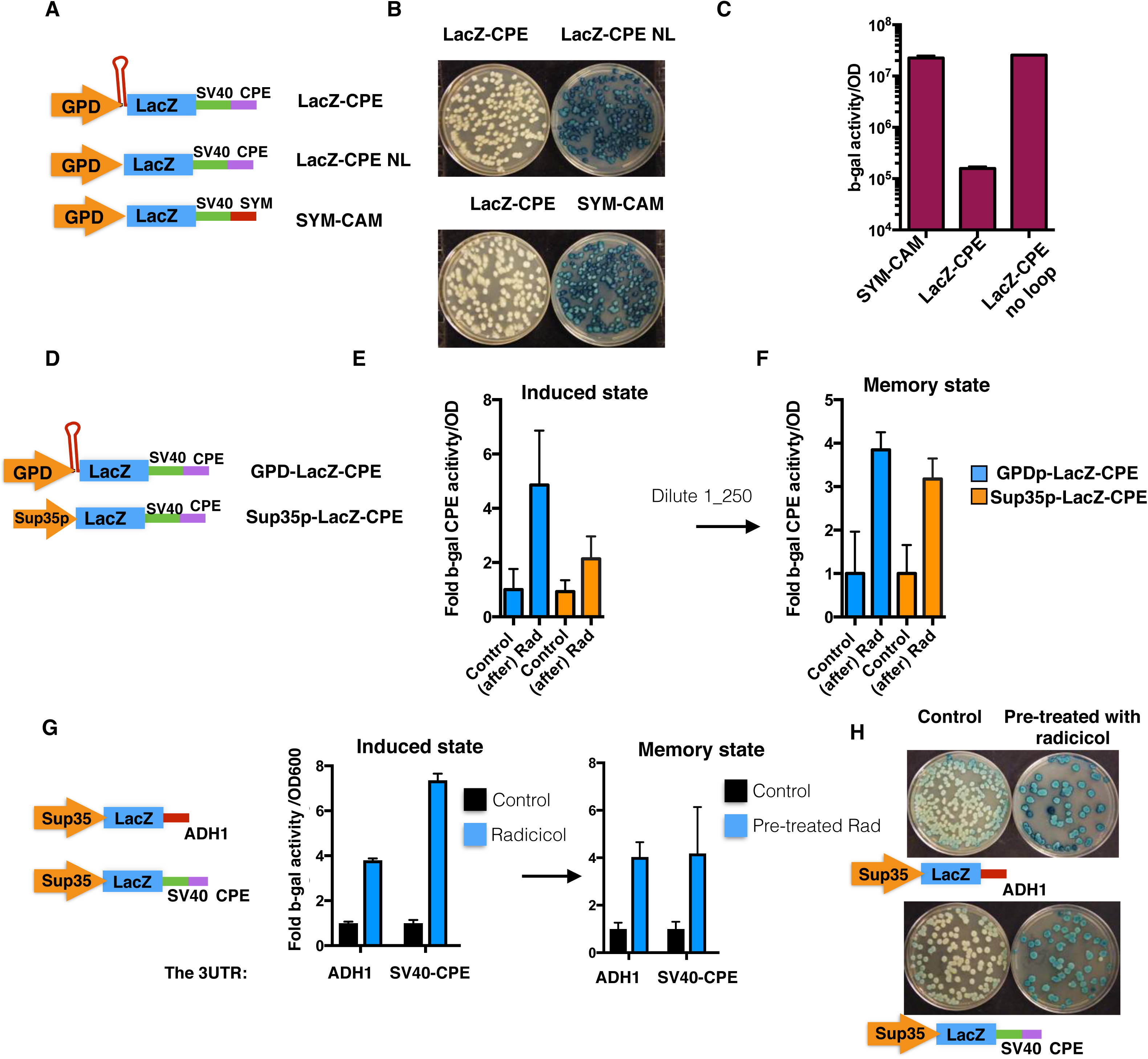
(A) Schematic of translation reporter constructs. LacZ-CPE is the original reporter used in previous studies (ref); LacZ-CPE NL is the same as LacZ-CPE but without a predicted stem loop in the 5’ UTR; and LacZ SYM-CAM is the original control from previous studies (ref). (B) Cells expressing the indicated translation reporters were grown in liquid culture, plated and then replica plated on x-gal plates to visualize the LacZ activity. (C) The activity of b-gal in strains harboring the indicated translation reporters was measured and plotted after overnight growth. (D) Schematic of the previously described translation reporter (ref) LacZ-CPE and the newly constructed translation reporter (Sup35p-LacZ-CPE) which lacks the stem loom at the 5’ UTR and has a weaker promoter (Sup35p). (E-F) comparison of radciciol activation (E, induced) and persistence (F, memory) (cells were grown over night and then diluted 1:250 and grown in the absence of radicicol) of translation of both the LacZ-CPE and the new Sup35p-LacZ-CPE reporter. (G) Schematic representation of the translation reporters with alternative 3UTR sequences (left). The relative induction of b-gal activity by radicicol was measured in cells harboring the distinct translation reporters (Induced state) and the degree of persistence following dilution 1:500 and overnight growth in the absence of the inhibitor (Memory state). (H) Cells harboring the indicated translation reporters were grown in the presence of 10uM radicicol ON and then plated on plates lacking radicicol. The plates were then replica plated onto x-gal plates and the relative b-gal activity was visualized.

**Sup figure 4.**
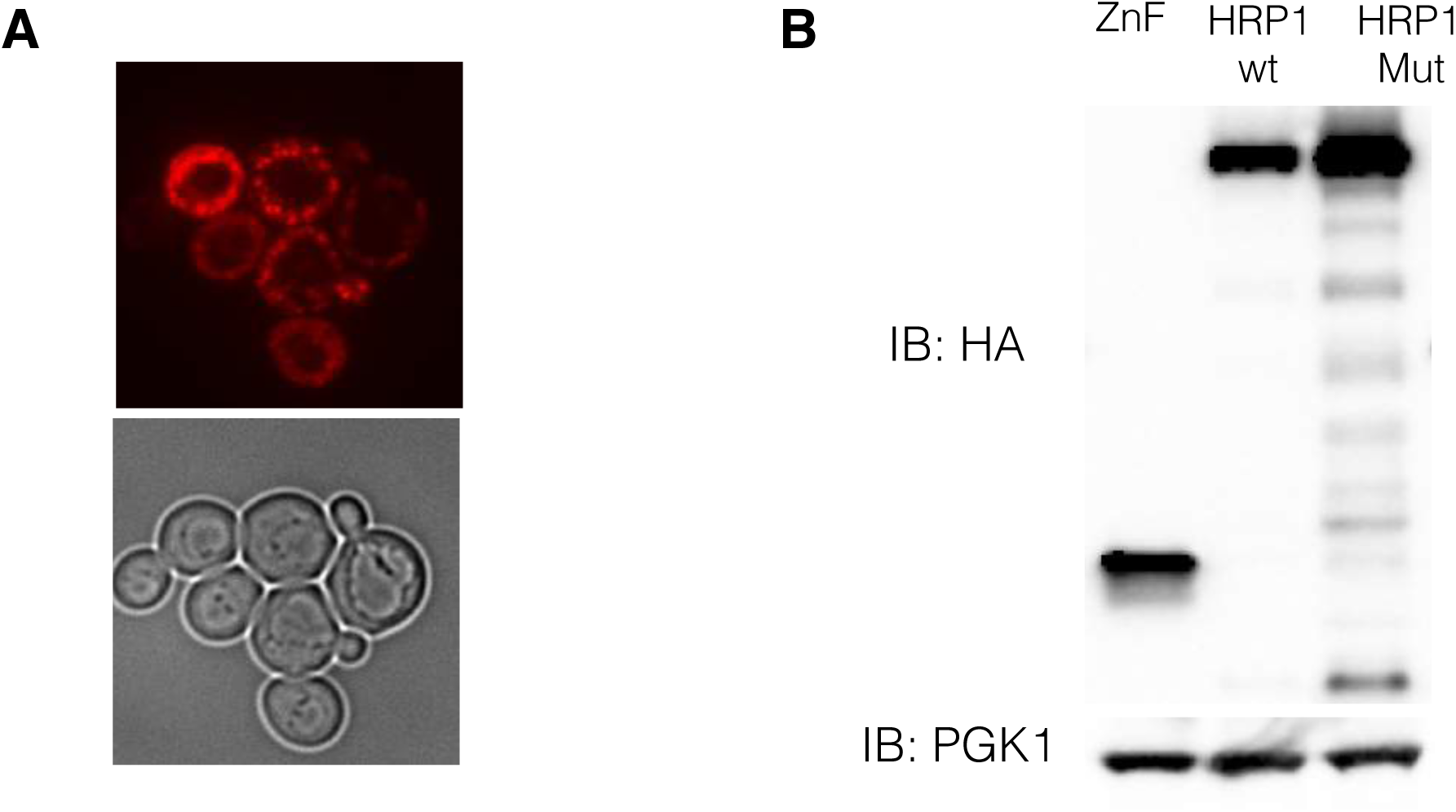
(A) Visualization of overexpression of the mkate tagged HRP1 PY521/2AA mutant after 6 hours of induction in galactose media. (B) Western blot analysis of total protein levels in cells overexpressing the wild type HRP1, PY521/2AA HRP1 mutant and ZnF control yTRAP sensors.

**Sup figure 5.**
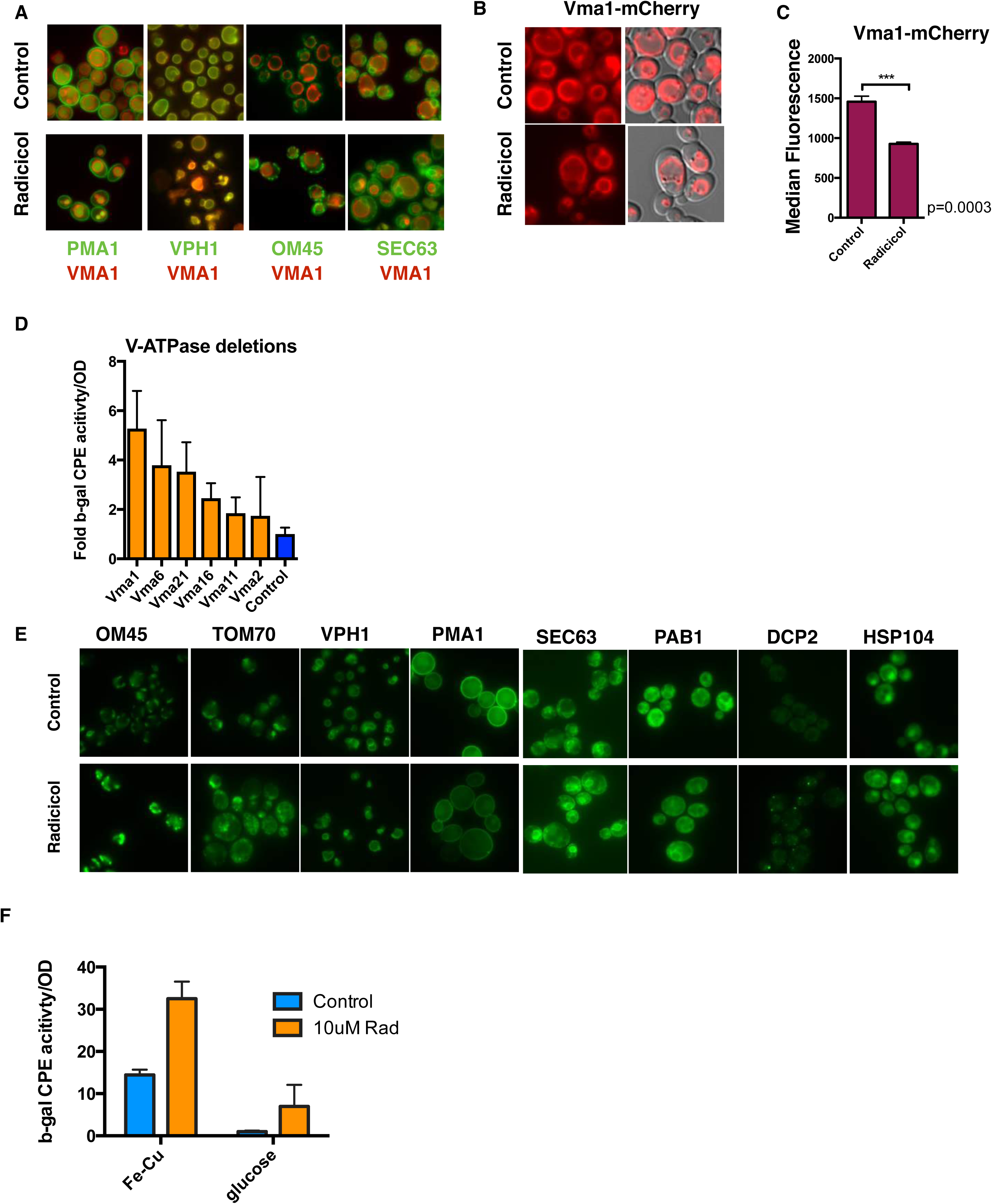
(A) Vma1 (tagged with mCherry) localization with respect to the vacuolar protein (VPH1), plasma membrane protein (PMA1), Mitochondrial protein (OM45) and ER-localized proteins (SEC63) all tagged with GFP was analyzed after overnight treatment with or without 10uM radicicol. (B-C) VMA1 endogenously tagged with mCherry is visualized (B) and levels quantified (C) following overnight growth with or without 10uM radicicol. *** p=0.0003. unpaired t-test analysis was used. (D) Strains with the indicated V-ATPase subunit deletions expressing the CPEB translation reporter were analyzed for their relative b-gal activity. (E) GFP-tagged OM45, TOM70, VPH1, PMA1, SEC63, PAB1, DCP2 and HSP104 were visualized following overnight treatment with or without 10uM radicicol. (F) LacZ activity in cells harboring the CPEB translation reporter grown overnight in normal (glucose) or iron-copper depleted media (-Fe-Cu) in the presence or absence of 10uM radicicol.

